# Persistence of Anti-Zika Virus Immunoglobulin M Antibodies in Children with Microcephaly up to Four Years after Primary Infection

**DOI:** 10.1101/857847

**Authors:** Gubio S. Campos, Rejane H. Carvalho, Maria da Glória Teixeira, Giovanna F. Britto e Silva, Carolina A. Rolo, Aline D. L. Menezes, Rafael. R. M. Souza, Silvia I. Sardi

## Abstract

Zika virus (ZIKV) is a member of the flaviviridae family of virus, considered to cause acute self-limited infection in adults, though it may lead to severe complications. It is believed that ZIKV infection elicit a classical viral immune reaction, with primary IgM antibody response and secondary IgG immunity. Persistence of IgM antibodies has been identified for other viruses belonging to the same family as ZIKV. We investigated, therefore, the presence of anti-ZIKV IgM antibodies in children with microcephaly born between January 2015 and November 2018, and their parents. We have detected persistence of IgM in 22% of children with microcephaly up to four years after primary infection. Long term IgM persistence have implications for the diagnosis of acute infection. More investigation is needed in order to correctly construe the significance of anti-ZIKV IgM persistence in the population in general, and in children with microcephaly in particular. The dynamics of IgM antibody responses against ZIKV must be known and understood to avoid misinterpretation of diagnosis for acute infection, re-infection and antibody persistence.

## Introduction

The Zika virus (ZIKV) belongs to the flavivirus genus, along with the Dengue virus (DENV), West Nile virus (WNV), Yellow Fever virus (YFV) and others. In most cases, infection by those viruses causes a self-limited illness, presenting, in the acute phase, general symptoms such as fever, myalgia, joint pain and rash. However, it has been already established that ZIKV can infect adult neurons and neural progenitor cell (1, 2), hence the disease may occasionally evolve to a neurological disorder. In adults it manifests itself as Guillain–Barré syndrome (3), while when acquired during pregnancy, ZIKV may affect the fetus neurological development and cause microcephaly in infants (4), known as congenital zika syndrome (CZS).

Despite disease severity, it is believed that the virus clears the body due to the immune response. Subjects with acute ZIKV infection presents a primary immune response characterized by the increase of anti-ZIKV IgM levels four to seven days after symptoms, which may be detected in body fluids, such as serum, saliva, urine and cerebrospinal fluid (5). In a classical viral immune response, levels in body fluids should decline about two to three months after symptoms onset, with the increase of anti-ZIKV immunoglobulin G (IgG), which will remain detectable, preventing reinfection.

Although cases of persistence of IgM in serum have been demonstrated for other Flaviviruses (6, 7), no study has investigated the levels of anti-ZIKV IgM for more than two years, nor there is a study conducted on long term IgM detection in children with microcephaly. Persistence of IgM antibodies may hinder diagnosis of acute infection in clinical settings, and interfere in serological surveillance in epidemiological studies. We are, therefore, interested in investigating whether ZIKV elicits IgM antibodies to persist in sera, and in following up for how long they remain detectable.

## Methods

### Study design and subjects

Our group is conducting a study on persistence of ZIKV and anti-ZIKV antibodies in children with CZS in the state of Bahia, Brazil. Circulation of ZIKV in Bahia was first described during an outbreak in 2015 (8). Later, ZIKV infection was associated with the development of microcephaly in children born to mothers infected by the virus during pregnancy (4).

The study population is composed of children with CZS, and their mothers, born from 2015 onwards. At the present, 37 children: eighteen males (49%) and nineteen females (51%), and their mothers (total n = 73) have been enrolled in the study. All the children were born between January 2015 and November 2018. Informed consent to participate in this study was obtained from the mothers, who also provided the informed consent on behalf of their child. This study was approved by the Ethical Committee of the Instituto de Ciências da Saúde –UFBA under the project name: Zikamarks, number 82340017.9.0000.5662.

### Anti-ZIKV IgM and Anti-DENV IgM assays

Blood samples were collected and processed by centrifugation for separation of serum. Serum samples were then divided into aliquots and froze at −80 □C until tests were performed.

We have tested the sera for the presence of anti-ZIKV IgM antibodies using the Vircell ZIKA ELISA IgM commercial kit. Considering that serology tests between ZIKV and DENV cross-react, we have also tested the anti-ZIKV IgM positive samples for the presence of anti-DENV IgM using the Panbio Dengue IgM Capture ELISA. All commercial testes were carried out according to manufactures protocols, and calculations for positive, negative and equivocal results were performed as per the manufacturer instruction.

## Results

We have detected that from the 37 children with microcephaly tested, 10 of them still presented high levels of anti-ZIKV IgM antibodies, and from those, only 2 were also positive for anti-DENV IgM (Table 1). Considering the possibility of cross-reaction between ZIKV and DENV antibodies, at least 8 children presented persistent anti-ZIKV IgM levels in serum. From those, 2 were male (25%) and 6 were female (75%). Chi-square analysis (0.5% confidence interval) did not show correlation between long term production of anti-ZIKV IgM antibodies and child gender.

**Table 1:** Data from the children and anti-zika virus IgM assays.

| Child | ZIKV | DENV | Birth |  |
| --- | --- | --- | --- | --- |
| ID | IgM | IgM | date | Gender |
| MC01 | NEG | NA | October 15 | F |
| MC02 | NEG | NA | November 15 | M |
| MC03 | NEG | NA | ND | M |
| MC04* | POS | NEG | November 15 | F |
| MC05 | NEG | NA | November 15 | M |
| MC06 | NEG | NA | ND | F |
| MC07 | NEG | NA | ND | M |
| MC08 | NEG | NA | November 15 | F |
| MC09 | NEG | NA | November 15 | F |
| MC10* | POS | NEG | ND | F |
| MC11 | NEG | NA | ND | F |
| MC12* | POS | NEG | July 15 | F |
| MC13 | NEG | NA | February 16 | F |
| MC14 | NEG | NA | December 15 | M |
| MC15 | POS | POS | December 15 | F |
| MC16 | NEG | NA | December 15 | M |
| MC17 | NEG | NA | May 16 | F |
| MC18 | NEG | NA | ND | M |
| MC19 | POS | POS | May 16 | F |
| MC20* | POS | NEG | ND | F |
| MC21* | POS | NA | ND | M |
| MC22 | NEG | NA | ND | M |
| MC23 | NEG | NA | June 15 | M |
| MC24 | NEG | NA | June 15 | M |
| MC25 | NEG | NEG | November 18 | M |
| MC26* | POS | NA | November 15 | F |
| MC27 | NEG | NA | January 16 | M |
| MC28 | NEG | NA | July 16 | M |
| MC29 | NEG | NA | Dec, 15 | M |
| MC30* | POS | NEG | November 15 | F |
| MC31* | POS | NEG | January 16 | M |
| MC32 | NEG | NA | January 16 | F |
| MC33 | NEG | NA | December 15 | F |
| MC34 | NEG | NA | July 15 | M |
| MC35 | NEG | NA | November 15 | M |
| MC36 | NEG | NA | October 15 | F |
| MC37 | NEG | NA | July 15 | F |

\* Cases with persistent IgM antibodies; POS = Positive; NEG
= Negative; NA = Not applicable; ND = information not disclosed; F = female; M = male.

## Discussion

We report the persistence of anti-ZIKV IgM antibodies in serum of children with microcephaly up to four years after symptoms onset and primary antibody response. Serology test for anti-DENV IgM showed negative, demonstrating there was no cross-reactivity.

Although it is well known that serological assays for ZIKV antibodies may cross react with other Flaviviruses besides DENV (9), there have been no outbreaks of other flaviviruses, but ZIKV and DENV in the region; there is no confirmation of WNV circulating in the state of Bahia, and epizootic YFV has been reported in monkeys (10), but not in humans.

The IgM production is the first line of humoral response after the presentation of antigens on lymph nodes. In general, the IgM production start in the very beginning of the viral infection, reaching higher levels about day 6, and starting to decrease until the third month. Prolonged IgM production could be related to the re-exposure of ZIKV antigens either from the environment or from viral deposited within specific tissues in the body, since some virus has the ability of evading the immune response, sneaking, and depositing in specific tissues (11). It is, therefore, possible that virus deposits in health individual’s bodies can stimulate a new primary humoral immune response maintaining IgM high levels.

Investigation of ZIKV deposits in tissues revealed that primates infected subcutaneously with ZIKV presented detectable viral RNA over 60 days in the central and peripheral neuronal tissues, and other deposits have been already described in the for Rhesus macaque and Red-bellied tamarin infected models (12).

Persistence of IgM after a flavivirus infection has been observed. Anti-WNV IgM could still be detected even after 8 year (7), and immune response to YFV vaccine arouse long term IgM presence (6). Recently, anti-ZIKV IgM and neutralizing antibodies were detected in sera up to 19 months after infection (13).

Although it was not significant, we have observed a slight tendency for IgM persistence in female children (25% male vs. 75% female), while the sample distribution between males and females were 49% and 51% respectively. Increasing the amount of subject in the study may resolve the tendency or definitely proof there is no dependence between those two variables.

## Conclusion

More investigation is required, more data needs to be produced and studies compared in order to correctly construe the significance of anti-ZIKV IgM persistence in the population in general, and in children with microcephaly in particular. And, despite the cause for long persistence of IgM regarding ZIKV infection being unknown, its implication for the diagnose of acute infection is clear. The dynamics of IgM antibody responses against ZIKV must be known and understood to avoid misinterpretation of diagnosis for acute infection, re-infection and antibody persistence.

## Acknowledgments

We are grateful to CAPES-ZIKA Fast Track, FAPESB APP0076/2016, and FINEP/CAPES ZIKA Biomarkers for the financial support.

